# Chromosome engineering points to the *cis*-acting mechanism of chromosome arm-specific telomere length setting and robustness of plant phenotype, chromatin structure and gene expression

**DOI:** 10.1101/2024.12.09.627491

**Authors:** Ondřej Helia, Barbora Matúšová, Kateřina Havlová, Anna Hýsková, Martin Lyčka, Natalja Beying, Holger Puchta, Jiří Fajkus, Miloslava Fojtová

## Abstract

The study investigates the impact of targeted chromosome engineering on telomere dynamics, chromatin structure, gene expression, and phenotypic stability in *Arabidopsis thaliana*. Using precise CRISPR/Cas-based engineering, reciprocal translocations of chromosome arms were introduced between non-homologous chromosomes. The subsequent homozygous generations of plants were assessed for phenotype, transcriptomic changes and chromatin modifications near translocation breakpoints, and telomere length maintenance. Phenotypically, translocated lines were indistinguishable from wild-type plants, as confirmed through morphological assessments and principal component analysis. Gene expression profiling detected minimal differential expression, with affected genes dispersed across the genome, indicating negligible transcriptional impact. Similarly, ChIPseq analysis showed no substantial alterations in the enrichment of key histone marks (H3K27me3, H3K4me1, H3K56ac) near junction sites or across the genome. Finally, bulk and arm-specific telomere lengths remained stable across multiple generations, except for minor variations in one translocation line. These findings highlight the remarkable genomic and phenotypic robustness of *A. thaliana* despite large-scale chromosomal rearrangements. The study offers insights into the *cis*-acting mechanisms underlying chromosome arm-specific telomere length setting and establishes the feasibility of chromosome engineering for studies of plant genome evolution and crop improvement strategies.

**Significance statement:** This study demonstrates the robustness of *Arabidopsis thaliana* in maintaining telomere stability, chromatin integrity, and wild-type phenotype despite large-scale chromosomal translocations. It underscores the potential of chromosome engineering in advancing genome evolution research and crop improvement, contributes to a deeper understanding of genome dynamics, and opens new avenues for precision breeding in plants.

## Introduction

Current approaches for targeted genome editing have significantly improved the accuracy and speed of generating organisms with modified genomes. Many of them have been rapidly adopted in plant research, including the improvement of crops. Among these methods, clustered regularly interspaced short palindromic repeats associated with the Cas nuclease (CRISPR/Cas) have emerged as a gold standard for inducing double-stranded DNA breaks (DSBs) at specific genomic sites in both basic and applied research. Utilization of multiple guide RNAs, which provide targeting specificity, enables the introduction of several DSBs. This advancement has expanded the potential of genome modifications – from simple single-gene editing to complex large-scale chromosome engineering and simultaneous targeting of multiple genes (Pan et al., 2022; Pan et al., 2021; Zhang et al., 2021). The high specificity of these genome-editing techniques, demonstrated by low off-target and other unpredictable activities (Bessoltane et al., 2022), holds great promise for their more extensive application in crop breeding.

Large-scale genome rearrangements have occurred naturally during plant evolution, contributing significantly to plant genome adaptation and speciation (reviewed in Schubert and Vu, 2016). These events can lead to various consequences: (i) the loss of function of previously active genes, (ii) changes in open reading frames, (iii) modulation of gene expression patterns, and (iv) shifts in functionally specialized chromosome regions (e.g., centromeres), ultimately disrupting existing genetic linkages or forming new ones (Fransz et al., 2016; Lowry and Willis, 2010; Schubert, 2018). The possibilities offered by CRISPR-based techniques have greatly advanced our capabilities for the directed evolution of plants (reviewed in Zhang et al., 2019), facilitating the ability to generate plants that can better cope with climate changes and other environmental stressors.

Generally, DSBs on heterologous chromosomes can introduce reciprocal translocations (Pacher et al., 2007), while DSBs on the same chromosome can result in deletions or inversions (Qi et al., 2013; Siebert and Puchta, 2002). Two DSBs were induced in the arms of *Arabidopsis thaliana* chromosomes 1 and 2 or chromosomes 1 and 5 (Beying et al., 2020) using a codon-optimized Cas9 nuclease from *Staphylococcus aureus*, which induces error-prone, efficient and stably inherited non-homologous end-joining-mediated mutagenesis (Steinert et al., 2015) in an egg cell-specific expression system (Wang et al., 2015). SaCas9s were targeted to intergenic regions approximately 0.5 or 1.0 Mb from the ends of long arms of respective chromosomes (Figure 1A), and homozygous lines with these large translocated regions were selected. This system offers exciting potential for breeding programs, as the targeted introduction of reciprocal translocations may lead to the selection of favorable quantitative traits. However, the long-term and complex consequences of such extensive chromosomal rearrangements require further exploration.

**Figure 1:**
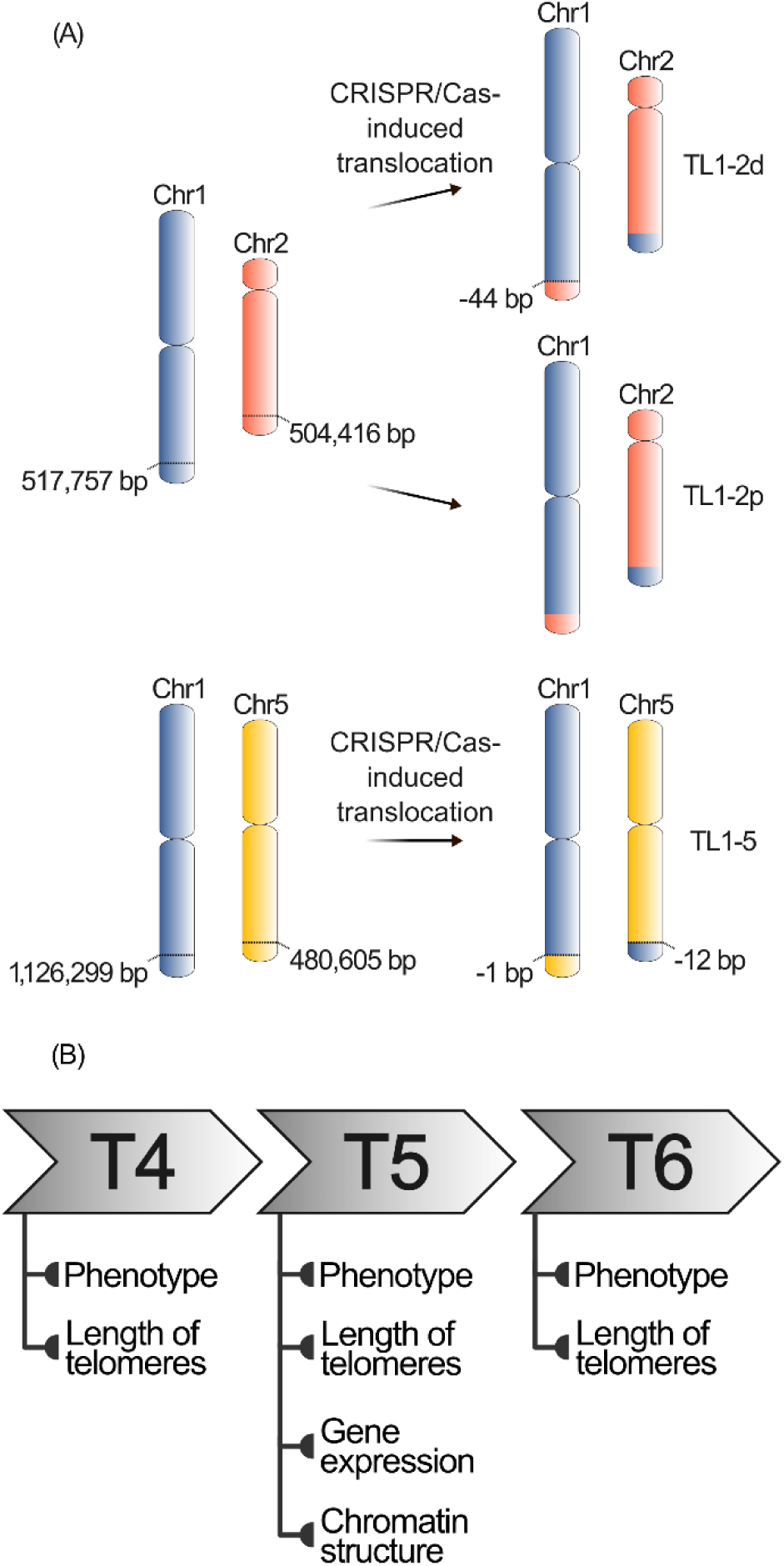
Description of *A. thaliana* lines with translocated chromosome arms and experimental design. (A) Construction and selection of *A. thaliana* lines with translocated chromosome arms are described in Beying, et al. (2020). By translocation of approximately 500 kb of long arms of chromosomes 1 and 2, the TL1-2d line (with 44 bp deletion at junction 1), and the TL1-2p line (with perfect ligation at both junctions), were constructed. By translocation of approximately 1 Mb and 500 kb of long arms of chromosomes 1 and 5, respectively, the TL1-5 line with small deletion at both junctions was constructed. (B) Representatives of three consecutive generations of plants with translocated chromosome arms (T4, T5 and T6, i.e., the second, third, and fourth homozygous generations following transformation) were analyzed for the length of telomeres and phenotype (T4, T5, T6), gene expression at the level of transcripts and chromatin structure (T5).

Telomeres are functionally important regions located at the translocated parts of chromosomes. Telomeres are nucleoprotein structures that cap the ends of linear eukaryotic chromosomes and play a crucial role in maintaining genome stability and cellular longevity. They prevent natural chromosome ends from being recognized as DSBs and protect them from inappropriate repair, thus addressing the “end-protection problem”. Furthermore, telomeres prevent the loss of the coding sequences caused by the progressive shortening of chromosome ends due to their incomplete replication, i.e. “end-replication problem”. Despite the critical importance of telomeres in maintaining genome integrity, many unanswered questions persist, including the regulation of telomere length and the factors that influence it. Telomere length is an extremely variable parameter across organisms. In plants, telomere lengths range from less than 0.5 kb in algae (Fulnecková et al., 2012) to more than 100 kb in tobacco (Fajkus et al., 1995). No obvious relationship has been found between telomere lengths and the genome size, number of chromosomes, and the sequence of telomere repeat (Adamusova et al., 2020). Considerable variations in telomere lengths have also been observed even among individuals of the same species. For instance, telomeres in inbred lines of maize range from 1.8 kb to 40 kb (Burr et al., 1992), and Arabidopsis ecotypes exhibit telomeres ranging from 2 kb to 9 kb (Shakirov and Shippen, 2004). Generally, the balance between their positive and negative regulators determines telomere lengths at the cellular level. Factors involved in the assembly of the telomerase holoenzyme – an enzyme complex capable of elongating telomeres – and its recruitment to chromosome ends (reviewed in Schmidt and Cech, 2015), as well as telomere-binding and telomere-associated proteins that limit telomerase access to telomeres (Fulneckova and Fajkus, 2000), exemplify this balance. However, the general cellular balance cannot explain the fact that even within a single cell, telomeres on different chromosome arms are maintained at distinct, yet relatively stable, lengths. The rationale behind this arm-specific telomere length regulation remains unknown, though *cis*-acting mechanisms, particularly the subtelomeric chromatin composition and structure, are thought to play a role. This hypothesis aligns with recent findings suggesting that the subtelomeric chromatin, particularly heterochromatin spreading, plays a role in setting the specific telomere length in budding yeast (Teplitz et al., 2024).

The current availability of plants with translocated chromosome ends offers a unique possibility to address the issue of chromosome-arm-specific telomere length setting. Here, we utilized previously generated plant lines in which telomeres at 1R, 2R and 5R chromosome arms have been spatially relocated due to the precise chromosome engineering events (Figure 1A). In plants with these chromosome arms translocations, we examined the impact of these chromosome structure changes on plant phenotype, gene expression and chromatin structure, particularly in regions adjacent to the translocation breakpoint (for experimental design, see Figure 1B). Next, bulk telomere lengths, as well as lengths of telomeres at specific chromosome arms, were analyzed. Our results demonstrate the robustness of the *A. thaliana*, which can tolerate these large-scale chromosome rearrangements over multiple generations.

## Results

### Plants with translocated chromosome arms retain wild-type phenotype

We monitored phenotypes of plants with translocated ends of R arms of chromosomes 1 and 2 with the small deletion at the J1 junction (TL1-2d); in plants with translocated ends of R arms of chromosomes 1 and 2 with perfect ligation at both junctions (TL1-2p); and in plants with translocated ends of R arms of chromosomes 1 and 5 (TL1-5; Figure 1A). Two different approaches were followed. Firstly, we did the standard photographing to document plants at the same developmental phases (5 weeks after sowing (5 WAS), and 12 WAS) as shown in Figure 2A. Plants with translocated chromosome arms maintained the WT phenotype during three consecutive homozygous generations.

**Figure 2:**
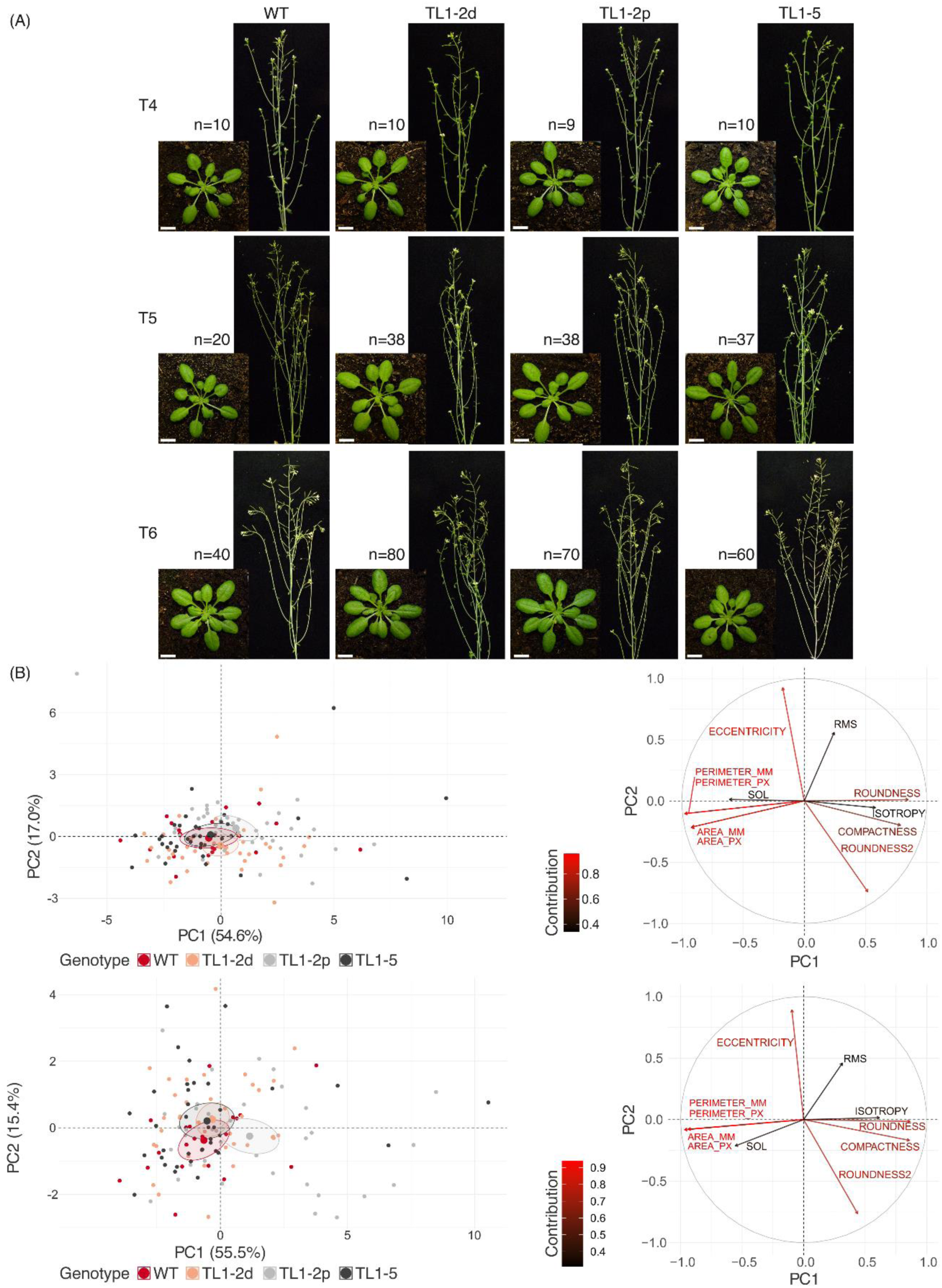
Phenotype of plants with translocated chromosome arms. (A) Photographs of WT plants and plants with translocated chromosome arms (T4, T5, T6 generations) at 5 WAS (left) and 12 WAS (right). (B) Visualization of RGB-imaging-based parameters as PCA plots (left) and bi-plots (right) of plants from T5 generation at 4 WAS (upper) and 5 WAS (lower); for T4 and T6 generations, see Figure S1. The length of the arrow (the longer the arrow, the higher the contribution) explains the contribution of tested variables. The differences in plant genotypes are indicated in PCA plots as centroids with ellipses representing their 95% confidence interval.

Next, we collected objective phenotypic data of 4 and 5 WAS plants for morphological parameters using the PlantScreen system (Photon Systems Instruments, Drásov, Czech Republic). The PlantScreen system can scan multiple morphological parameters of plants, from a raw area or perimeter to more complex ones like compactness or slenderness of leaves (11 parameters in total; (Pavicic et al., 2017)). Phenotypic data were evaluated by principal components analysis (PCA) for each age and generation of plants with translocated chromosome arms. For plants of T5 generation at 4 WAS, the first two principal component (PC) values explained 54.6% and 17.0% of the total variation, respectively. At 5 WAS, 55.5% and 15.4% of the total variation was explained by PCs. The overlap between ellipses of all three generations and both ages of plants depicted clear indistinguishability between plants of WT genotype and the ones with translocated chromosome arms at 4 WAS and slightly less so at 5 WAS (Figure 2B left; see Figure S1 for PCs values for T4 and T6 generations). PCA bi-plots showed only slight changes in the direction of variables between 4 and 5 WAS (Figure 2B right; see Figure S1 for the contribution of variables for T4 and T6 generations).

### Large-scale chromosome translocations do not affect transcriptome or chromatin structure

Genome editing techniques aimed primarily at modulating the expression of specific gene(s) through error-prone non-homologous end-joining repair of induced DSBs (Weeks et al., 2016). However, chromosome rearrangements affect 2D and 3D chromatin organization, which may subsequently dysregulate gene expression near junction sites. To investigate this, we compared the transcriptome and chromatin structure of WT plants to those with translocated ends of chromosome arms.

In the T5 generation of TL1-2d plants, 15 downregulated and 96 upregulated genes were detected. In TL1-2p plants, we observed 6 downregulated and 18 upregulated genes. TL1-5 plants showed minimal differential expression, with only three genes significantly upregulated (Figure 3, Table S1). Given the total number of detected transcripts (11 605), these differentially expressed genes (DEGs) constituted a minor fraction, with 0.96%, 0.21% and 0.03% of the total transcripts in TL1-2d, TL1-2p and TL1-5, respectively. Among those DEGs, 43, 13, and 3 were located on chromosomes with translocated arms in TL1-2d, TL1-2p and TL1-5, respectively (Table S1). Only one DEG in TL1-2d was located in the ± 100 kb window, and two DEGs in TL1-2d and one DEG in TL1-2p in the ± 500 kb window from the junction sites. This observation indicated that the majority of genes in these regions had stable transcript levels.

**Figure 3:**
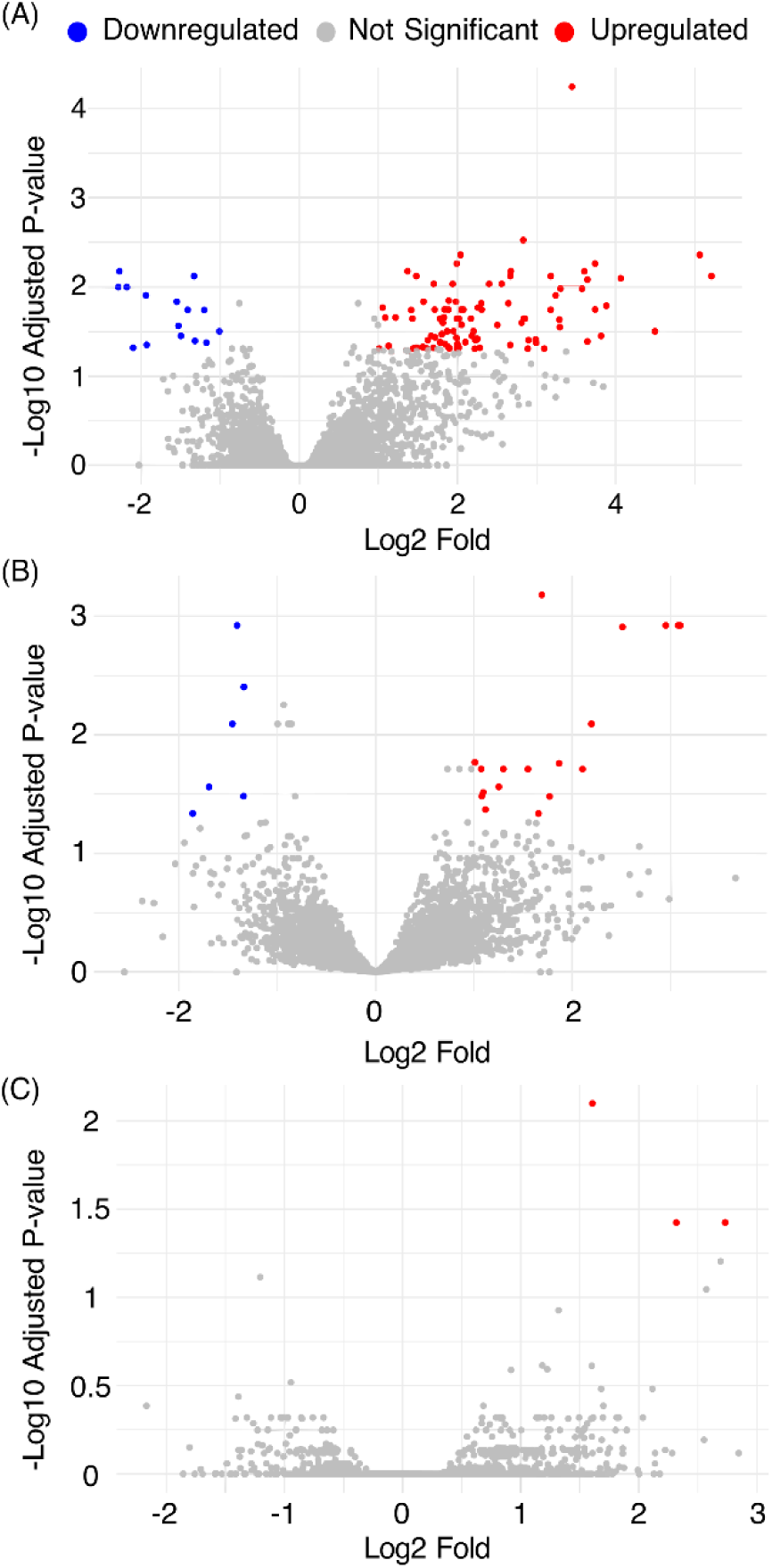
Volcano plots demonstrating DEGs in plant lines with translocated chromosome arms TL1-2d (A), TL1-2p (B) and TL1-5 (C). For RNAseq, 4 - 7 T5 plants of respective genotypes were selected (Table S3). For a description of DEGs and their positions on chromosomes, see Table S1.

To compare potential epigenetic changes due to the translocation of chromosome arms, we analyzed the enrichment of histone marks between the T5 generation of WT plants and plants with translocated chromosome arms by ChIPseq. We selected three histone modifications enriched on WT chromosomes near junction sites: Polycomb repressive H3K27me3 mark associated with developmentally silenced genes, and H3K4me1 and H3K56ac, which are linked to open chromatin regions. Enrichment of histone marks was analyzed in the T5 generation of plants with translocated chromosome arms. At the whole-chromosome level, the distributions of these histone marks were consistent, considering the translocation of chromosome ends (Figure 4). Analysis of regionś flanking sites of chromosome junctions (± 250 kb window) revealed no substantial changes in the enrichments of histone marks compared to the WT patterns (Figure 5).

**Figure 4:**
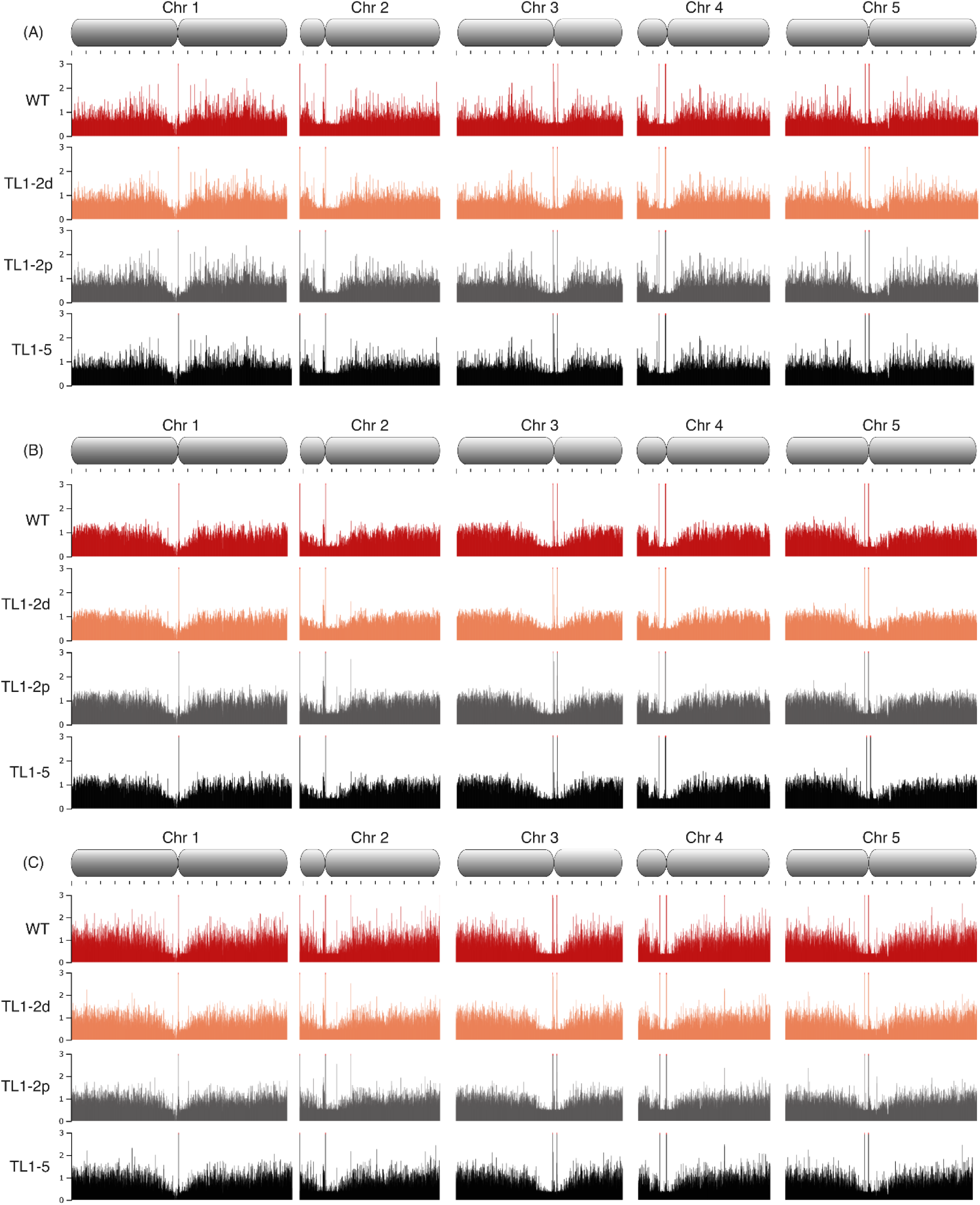
Profiles of histone epigenetic marks – H3K27me3 (A), H3K4me1 (B) and H3K56ac (C) across all five chromosomes of WT (red), TL1-2d (orange), TL1-2p (gray) and TL1-5 (black) plants. Visualization of ChIPseq data was done through JBrowse platform (Buels et al., 2016).

**Figure 5:**
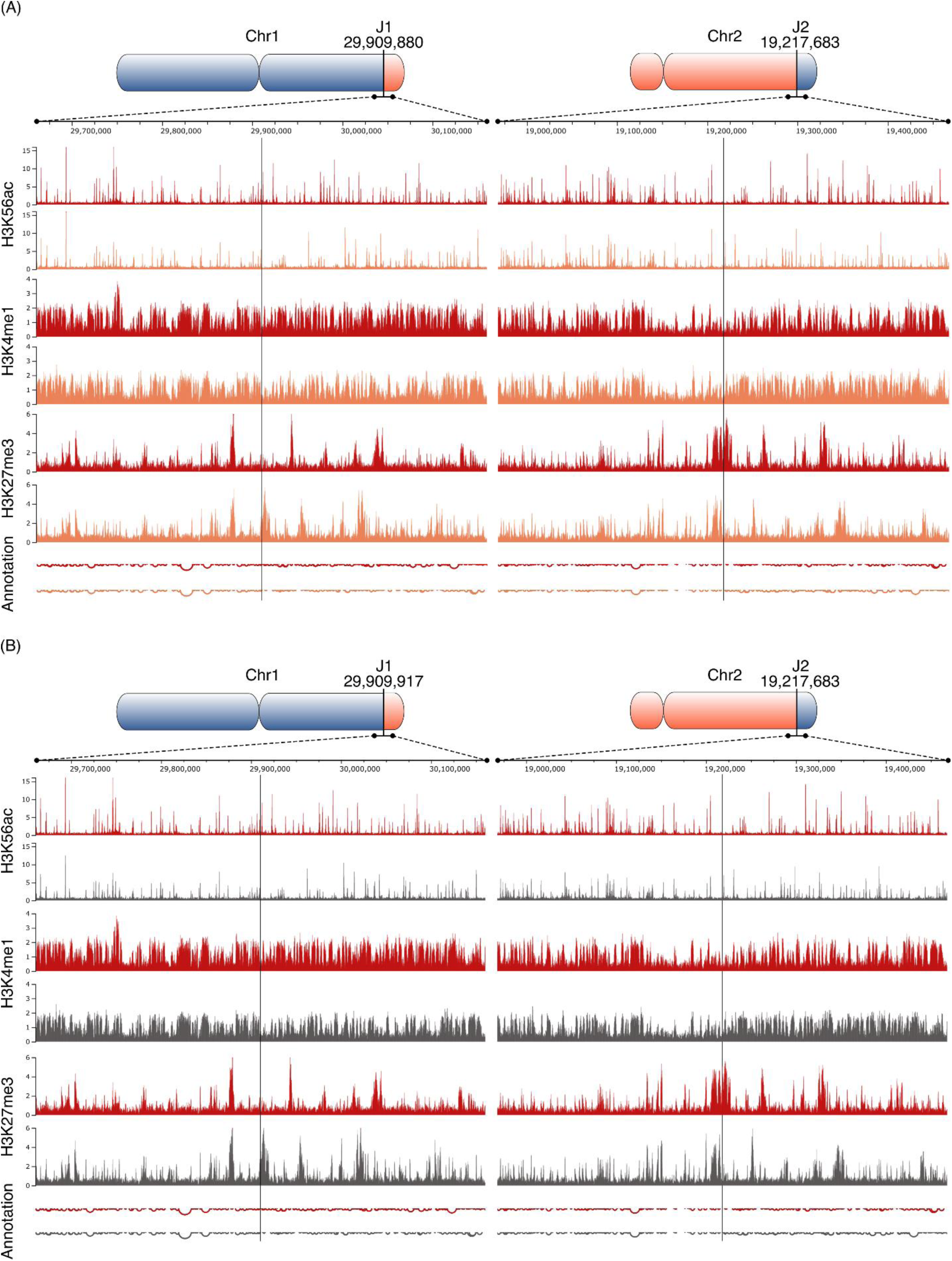
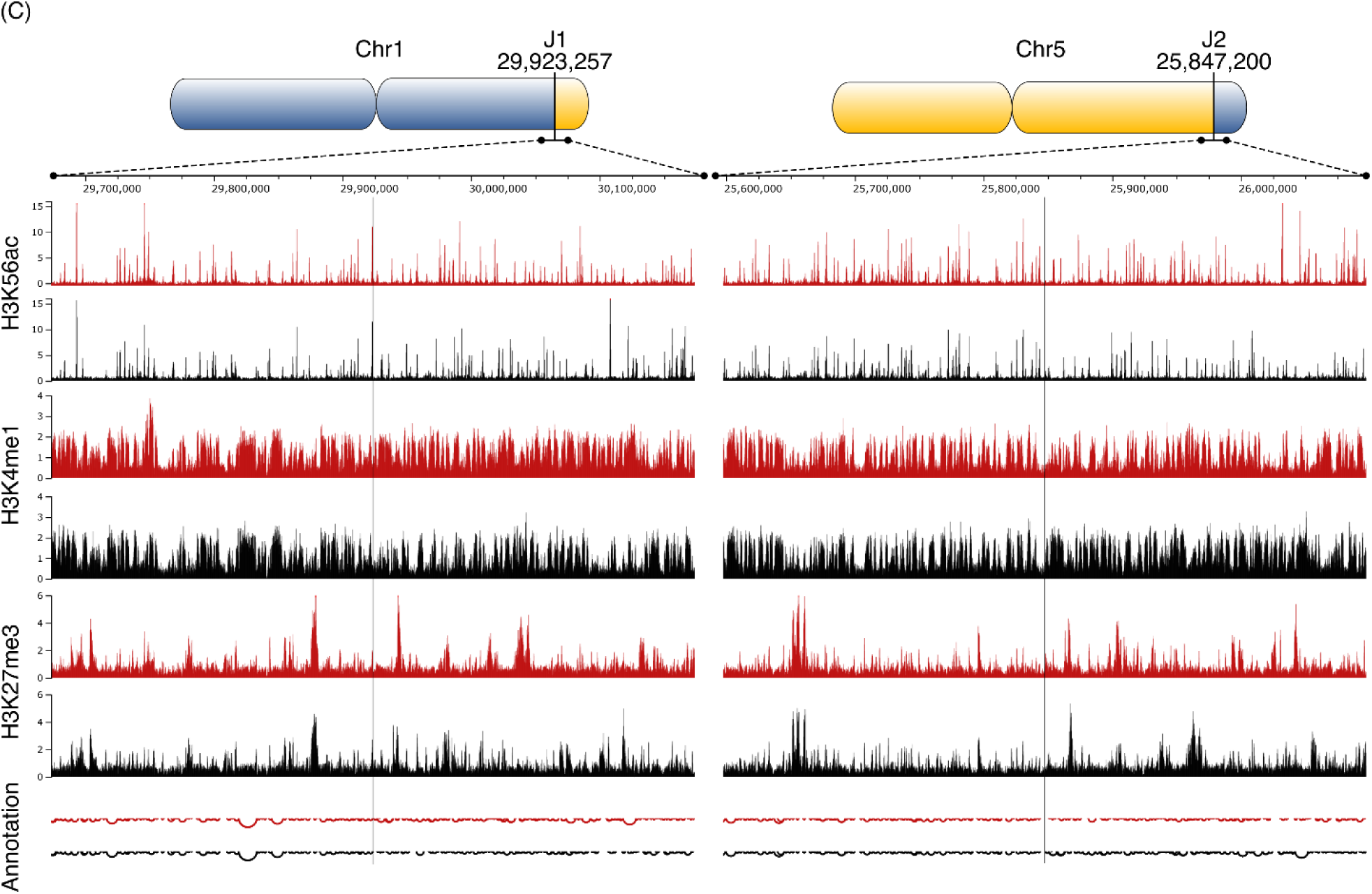
Detailed view of the enrichment of epigenetic marks in the window + 250 kb around junction sites in TL1-2d (A), TL1-2p (B), and TL1-5 (C). WT (red), TL1-2d (orange), TL1-2p (grey), TL1-5 (black). Annotation, depths of pits reflect the gene density in respective regions. Visualization of ChIPseq data wa done through the JBrowse platform (Buels et al., 2016).

In addition, the differential binding analysis was carried out to identify regions of the plant genome with statistically different enrichment of measured epigenetic histone marks. Only 48 differentially enriched regions were identified across all possible comparisons (Table S2). The highest number, 39 differentially enriched areas, was found in the comparison of WT vs. TL1-2d in the case of the H3K27me3 mark. However, only 16 of them were located on the chromosomes where the translocation occurred, and only 2 were in the ± 2 Mb window of the translocation point. Notably, none of the genes settled in these differentially enriched regions were found among DEGs (Table S1), rejecting the usually direct interconnection between chromatin structure and transcription. These findings show that extensive translocations of chromosome ends did not affect gene expression nor altered chromatin structure at the whole genome or chromosome level or near the translocation junctions.

### Telomeres are stable in plants with translocated chromosome ends across three consecutive generations

In *A. thaliana* of the Columbia ecotype, telomere lengths range approximately between 2.5 and 3.5 kb. Although the translocated regions are over two orders of magnitude longer (Figure 1A), the telomeres were shifted to the new chromatin environment in different chromosome territories, which may influence the telomere length setting. Bulk telomere length, i.e. lengths of telomeres across all chromosome arms, was analyzed by the terminal restriction fragment (TRF) method. Given the natural inter-individual variability in telomere lengths observed even in WT plants (Figures S2A, S3A), at least 25 individuals per translocated line across all three consecutive generations were analyzed to ensure reliable data (for numbers of analyzed plants, see Table S3). As is evident from the evaluation of telomere-specific hybridization signals, lengths of telomeres were preserved across all three consecutive generations of plants with translocated chromosome arms (Figures 6, S2B-D, S3A).

**Figure 6:**
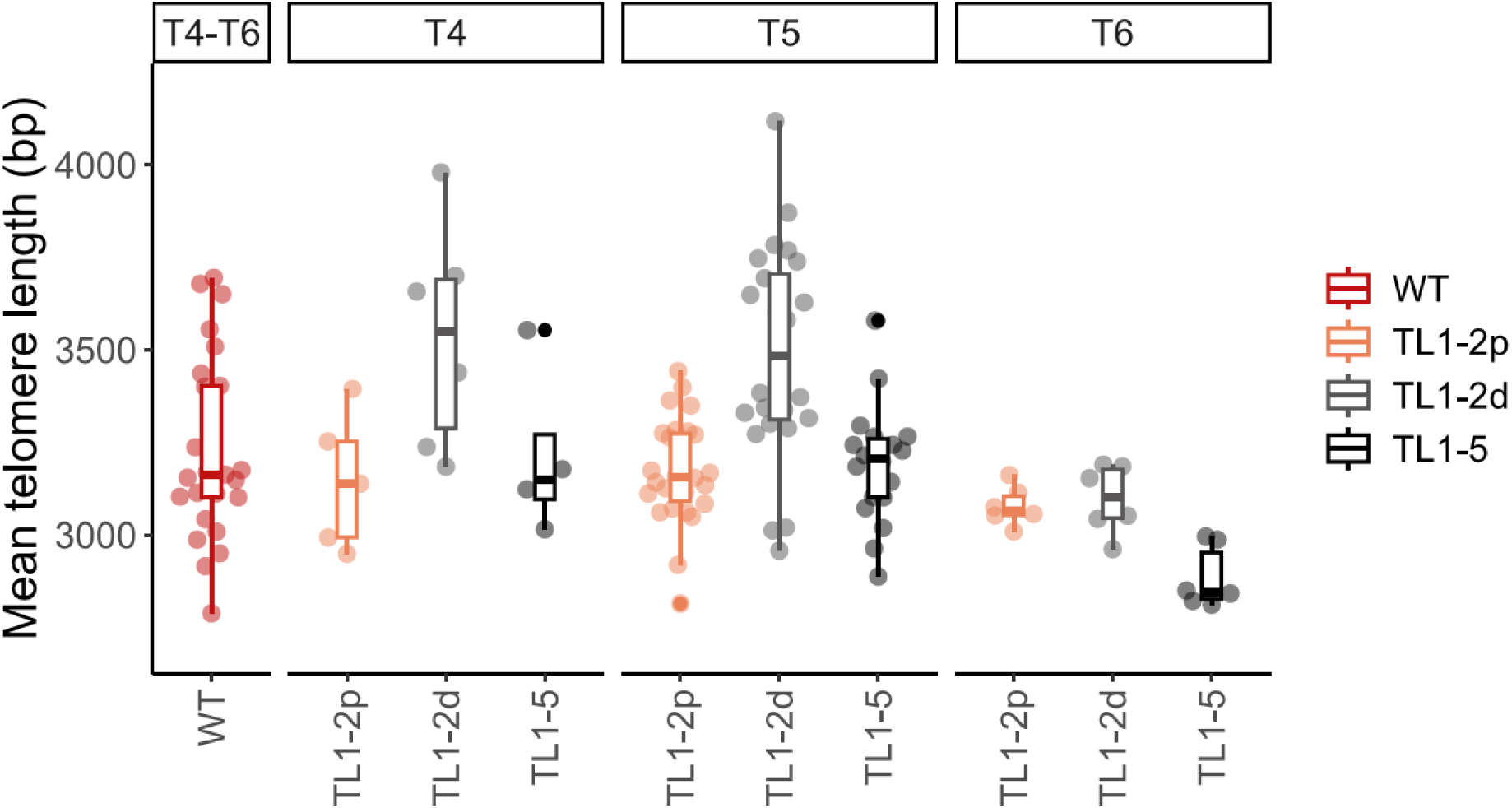
Summary of the TRF (terminal restriction fragments) analysis of bulk telomere lengths in plants with translocated chromosome arms of T4, T5 and T6 generations. Points represent the mean telomere lengths calculated based on the profiles of telomere-specific hybridization signals using the WALTER toolset (Lycka et al., 2021). Telomere lengths between combined generations of WTs and generations of mutants were compared by two-tailed Welch’s t-test according to the WALTER algorithm. For examples of raw hybridization data (WTs and T5 of plants with translocated chromosome arms) and WALTER outputs (T4, T5, T6), see Figure S2 and Figure S3A, respectively. For the number of plants analyzed, see Table S3.

To assess telomere lengths at translocated ends of specific chromosomes, we employed primer extension telomere repeat amplification (PETRA) protocol. This method leverages unique sequences in subtelomeric regions adjacent to each telomere, enabling amplification of the telomere on a specific chromosome arm (Heacock et al., 2004). Analysis of telomeres on 1R, 2R and 5R chromosome arms across all three generations of TL1-2p and TL1-5 plants showed that these large-scale chromosome rearrangements did not disrupt telomere homeostasis at translocated chromosome arms (Figures 7, S3B). Representatives of the TL1-2d line displayed shorter 1R telomeres in all generations and shorter 2R telomeres in T4 (Figures 7A&B, S3B).

**Figure 7:**
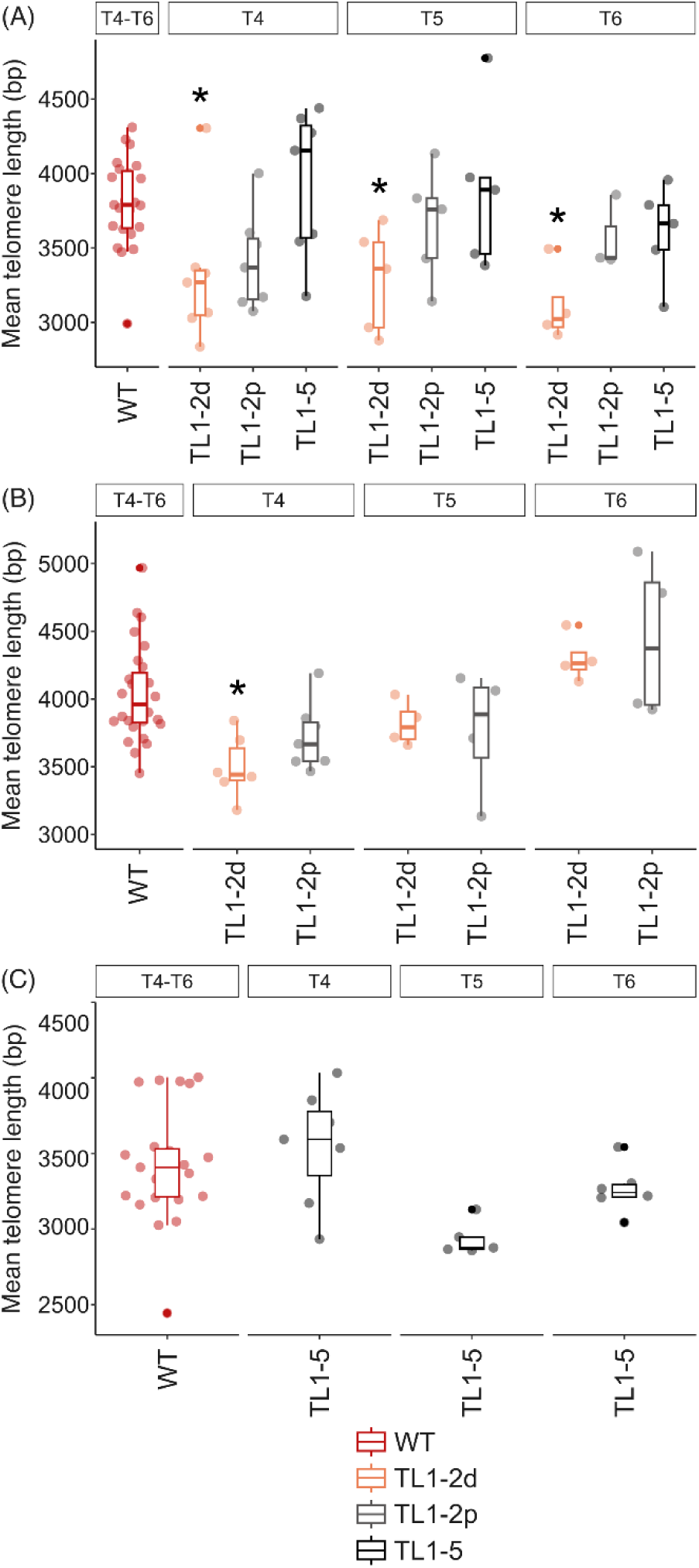
Summary of the PETRA (primer extension telomeric repeat amplification) analysis of telomeres located on 1R (A), 2R (B) and 5R (C) chromosome arms in plants with translocated chromosome ends of T4, T5 and T6 generations. Points represent the mean telomere lengths calculated based on the profiles of telomere-specific hybridization signals using the WALTER toolset (Lycka et al., 2021). Telomere lengths between combined generations of WTs and generations of mutants were compared by two-tailed Welch’s t-test according to the WALTER algorithm, * < 0.1 p-value (Lycka et al., 2021). For examples of raw hybridization data (WTs and T5 of plants with translocated chromosome arms) and WALTER outputs (T4, T5, T6), see Figure S4 and Figure S3B, respectively. For the number of plants analyzed, see Table S3.

## Discussion

Looking at plant evolution, there have been multiple instances where not only simple point mutations but also large-scale genome rearrangements have occurred (Schubert and Vu, 2016)). These spatial changes in chromatin structure are considered significant driving forces of genome evolution at the chromosomal level, as they have profoundly contributed to plant genome adaptation and speciation (Alkan et al., 2011; Schubert and Vu, 2016). One prominent example is reciprocal translocation, a genetic phenomenon involving the exchange of chromosome arms or segments between two non-homologous chromosomes. This process rearranges the genome without the loss of genetic information, classifying it as a balanced translocation. Reciprocal translocations play a crucial role in reducing chromosome numbers, often in conjunction with inversions and chromosome fusions (Lysak et al., 2006). In the context of plant genome evolution, reciprocal translocations are integral to fusion-fission cycles. During these events, telocentric chromosomes undergo translocations with breakpoints near centromeres or telomeres. Such rearrangements facilitate the transfer of chromosome arms, often leading to structural and functional modifications in the genome (Schubert and Vu, 2016).

The current karyotype of *A. thaliana* is thought to have evolved from the ancestral karyotype approximately 10 million years ago through two reciprocal translocations, three chromosome fusions, and at least three inversions (Lysak et al., 2006). Despite extensive studies, the detailed evolutionary events shaping the genome structure of *A. thaliana* remain unresolved, as well as the direct effects of these translocations and other chromosomal rearrangements on the plant genome. Advances in methodologies, including CRISPR/Cas-based chromosome engineering, now enable a precise real-time analysis of these evolutionary events; beginning with the immediate generations following the rearrangement (Khosravi et al., 2024). Beying et al. (2020) demonstrated that CRISPR/Cas could induce heritable reciprocal chromosome arm translocations mimicking natural evolutionary processes. In that study, translocations were successfully induced between chromosomes 1 and 2, and 1 and 5. Based upon this work, we investigated the immediate genomic responses to reciprocal translocations in subsequent generations, focusing on plant morphology, transcriptional activity, distribution of epigenetic histone marks, and telomere length maintenance.

To characterize plant phenotypes, we utilized both subjective (photography) and objective (phenotyping station) approaches. Across three subsequent generations of plants with translocated chromosome arms, neither photographs nor PCA (Figure 2) revealed distinguishable differences between WT plants and plants with chromosome arm translocations.

Reciprocal translocations can significantly affect gene expression, particularly when junction sites disrupt protein-coding genes or regulatory elements (Harewood and Fraser, 2014). Gene expression may also be impacted if translocation alters *cis*-regulatory elements up to 1.5 Mb from the junction site (Benko et al., 2009; Harewood and Fraser, 2014). In our study, most translocation junctions were located in intergenic regions, except for the pseudogene at J1 in TL1-5. The RNAseq analysis identified only a small number of DEGs (3-111 out of 11 605 transcripts; Table S1). These DEGs were dispersed throughout the genome, with only seven located within 1.5 Mb of the junction site (Table S1), suggesting the minimal impact of the translocations on gene expression.

Gene expression is closely linked to the chromatin structure. We analyzed the distribution of repressive (H3K27me3) and active (H3K4me1, H3K56ac) histone marks across entire chromosomes and in regions flanking translocation breakpoints. The ChIPseq analysis revealed no significant changes in the distribution of histone marks in translocated plants compared to WT, either at whole-chromosome level (Figure 4) or within ± 250 kb of junction sites (Figure 5). Importantly, the differential binding analysis identified only a few differentially enriched regions that were scattered throughout the genome (Table S2), with no corresponding changes in gene expression (Table S1). These findings suggest that chromatin structure and gene expression are robust to large-scale chromosomal translocations.

In general, the length of a telomere located at the end of each chromosome arm within the plant genome is determined by a complex interplay of genetic predispositions, environmental influences and intrinsic cellular mechanisms. While the cellular balance between positive and negative regulatory factors can explain the species-specific or ecotype-specific determination of telomere lengths, it does not account for the diversity and specific length settings of telomeres on individual chromosome arms. It is highly probable that *cis*-acting mechanisms play a role in this regulation. This hypothesis was supported by a recent study on yeast telomeres, which demonstrated that an increased abundance of the Sir4 protein in subtelomeric heterochromatin at a single chromosome end resulted in the lengthening of only that particular telomere (Teplitz et al., 2024). This is consistent with the role of Sir4 in recruiting telomerase mediated by the interaction with Ku80 protein, which binds the RNA component of yeast telomerase. However, the regulatory mechanisms in budding yeast may not directly apply to plant telomeres. For instance, the interaction between Ku80 and telomerase RNA has not been described in plants. Nevertheless, the core *cis*-acting principle of individual telomere regulation might be conserved across species. We hypothesized that chromosome arm translocations might affect telomere length homeostasis due to altered 3D chromatin structure. To test this hypothesis, bulk telomere lengths, as well as the lengths of specific chromosome-arm telomeres, were measured across three generations of plants. Results showed no significant differences between WT and translocated plants (Figures 6, 7), except for shorter telomeres on 1R (T4–T6) and 2R (T4 only) in TL1-2d plants. This may be attributed to a 44 bp deletion at the junction site in TL1-2d, which was absent in TL1-2p plants with the same translocation. Nevertheless, it is necessary to stress the variability of telomere lengths in WT plants as well as in plants with translocated chromosome arms (Figures S2, S3). Based on the results from telomere length analyses, the translocated parts of chromosomes may be too big to determine the *cis*-regulatory effect on telomere length, as telomeres form only minor parts of these regions (3 kb vs. 500 kb).

Our findings demonstrated that reciprocal translocations involving large chromosome segments likely include the transfer of regulatory regions maintaining the stability of telomere lengths and chromatin structure. Minor changes in gene expression and histone mark distribution appeared to reflect the natural variability rather than the direct effect of translocations. These results underscore the resilience of plant genomes to large-scale chromosomal rearrangements, providing insights into the mechanisms underlying genome evolution and stability.

The results presented in this article are also highly relevant for future breeding applications. Whereas by gene editing mainly individual traits can be agronomically improved, chromosome engineering gives us a tool at hand to control the combination of traits (Rönspies et al., 2021). Using Cas9, it has been demonstrated for plants in the last years not only that by reversing natural inversions recombination dead regions can be reactivated (Schmidt et al., 2020) but also that by the induction of novel inversions almost complete chromosomes can be excluded from genetic exchange (Rönspies et al., 2022). By the induction of reciprocal translocations between chromosome arms not only the linkage between genetic traits can be broken (Beying et al., 2020), in future we will also be able to create minichromosomes or even change chromosome numbers (Puchta and Houben, 2024). However, CRISPR/Cas-induced chromosomal rearrangements might also lead to unwanted secondary effects that could be detrimental to future breeding applications. Our study demonstrates that this is not the case – at least not for the analyzed translocations. No significant changes in phenotype, transcriptome, epigenome and telomere structure were found between engineered and wild-type plants making chromosome engineering a promising technology for future breeding applications.

## Experimental procedures

### Plant material

Homozygous lines of *Arabidopsis thaliana* plants of the Columbia ecotype with reciprocally translocated ends of long arms of chromosomes 1 and 2 (TL1-2d, 44 bp deletion at the junction site 1; TL1-2p, perfect ligation at both junction sites) or 1 and 5 (TL1-5) were described in Beying et al. (2020); see Figure 1A for the positions of translocations. Primers for genotyping are listed in Table S4, combinations of primers are described in Beying et al. (2020). Seeds were ethanol-sterilized and germinated in phytotrons for nine days on ½ Murashige and Skoog medium (Duchefa, Biochemical, Haarlem, The Netherlands; M0255.0050) supplemented with 0.8% agar under long-day conditions (16 h light, 100 mmol m^−2^s^−1^, 21 °C; 8 h dark, 19 °C). Seedlings were transferred to the soil and grown for five weeks under short-day conditions (8 h light and 16 h dark) to support leaf growth and then cultivated under long-day conditions to promote flowering. At 8 WAS, leaves were harvested for analyses of telomere lengths, RNAseq and ChIPseq.

### Analysis of plant phenotype

Plant phenotypes were monitored by photography in plants at 5 and 12 WAS. Next, the PlantScreenTM Compact System developed by Photon Systems Instruments (Drásov, Czech Republic), available at the Plant Sciences Core Facility (CEITEC, Masaryk University), was utilized. For each generation and line, 4 WAS and 5 WAS plants were scanned. The obtained images underwent pre-processing using the PlantScreen^TM^ Data Analyzer software to extract RGB-imaging-based parameters. Obtained morphological data were visualized and analyzed through PCA using the R package *factoextra* (Kassambara and Mundt, 2020). Additionally, the differences in plant genotypes were indicated as centroids with ellipses representing their 95% confidence interval.

### RNA isolation and RNAseq

Total RNA was isolated using NucleoSpin RNA Plant Mini kit (Macherey-Nagel) followed by DNaseI (TURBO DNA-free; Thermo Fisher Scientific) treatment. RNA quantity and quality were determined by Qubit4 fluorometer (Invitrogen) and agarose gel electrophoresis, respectively. The cDNA library for sequencing was prepared using QuantSeq™ 3’ mRNASeq Library Prep Kit FWD with UDIs (Lexogen) according to the protocol provided by the manufacturer. The cDNA library underwent sequencing using the Illumina NovaSeq 6000 platform with single-end read sequencing, resulting in a read length of 75 bp. Four to seven biological replicates were analyzed for each plant line (detailed in Table S3). The raw data are available in the NCBI SRA database under accession number PRJNA1195110.

### Differential gene expression analysis

Single-end 75 bp reads were aligned to the Arabidopsis TAIR10 genome (version 58, downloaded from plants.ensemble.org). The quality of the sequencing data was checked using FastQC (Andrews, 2010) and MultiQC (Ewels et al., 2016). Pre-processing of raw reads was done using Trimmomatic v0.38 (Bolger et al., 2014) with the following settings: SLIDINGWINDOW:4:20, HEADCROP:12, MINLEN:35. Next, pre-processed reads were mapped to the reference genome and transcriptome using STAR v2.7.10b (Dobin et al., 2013). Gene counts were quantified with RSEM v1.3.1. (Li and Dewey, 2011). Differential gene expression was analyzed using DESeq2 (Love et al., 2014). Genes with p-value <0.05 (corrected for multiple testing by the Benjamini-Hochberg method (Benjamini and Hochberg, 1995)), and log2 fold change values at least ±1 (i.e., fold change of the transcript level 2 and more, or 0.5 and less) were considered as DEGs.

### Native ChIPseq

Leaves were washed with sterile water and cross-linked with 1% formaldehyde (Sigma) for 10 minutes at room temperature under vacuum. The cross-linking process was quenched by glycine (final concentration 0.125 mM). The cross-linked leaves were stored at -80 °C.

Chromatin was isolated from 0.9 g of cross-linked leaves, following the procedure described by Saleh et al. (2008), up to obtaining nuclei pellets. From the step of nuclei lysis, the process continued as described by Vimont et al. (2020), using the MNase digestion for chromatin fragmentation. The chromatin was immunoprecipitated with the antibodies against H3K27me3 (ABE44, Millipore, USA), H3K56ac (07-677-I, Millipore, USA), and H3K4me1 (ab8895, Abcam, UK). DNA isolated from the input (chromatin without immunoprecipitation) and immunoprecipitated fractions (> 5ng per sample) was checked by the 5200 Fragment Analyser system (Agilent). Sequencing libraries from the DNA captured with chromatin immunoprecipitation were prepared using the Watchmaker DNA library prep kit (Watchmaker, USA), according to the manufacturer’s instructions. Adapters containing UMI sequence (xGen™ CS Adapters, IDT, USA) to enable detection of PCR duplicates were used. Libraries were PCR amplified for 6-14 cycles, depending on the input. The pooled library was sequenced on the MGI G400 instrument using DNBSEQ FCL PE200 cartridge, generating an average 10M reads per sequencing library. Three biological replicates were analyzed for all plant genotypes and histone modifications. The raw data are available in the NCBI SRA database under accession number PRJNA1195110.

### ChIPseq data evaluation

The quality of the sequencing data was assessed using FastQC (Andrews, 2010) and MultiQC (Ewels et al., 2016). The Illumina adapters and quality trimming of raw FastQC reads were performed using Cutadapt v4.3 (http://dx.doi.org/10.14806/ej.17.1.200) with settings: quality-base = 33; q = 0,20; m = 35; M = 250; a = AGATCGGAAGAGCACACGTCT; A = AGATCGGAAGAGCGTCGTGTA. The paired-end 95 bp reads were aligned using Bowtie2 v2.4.2 (Langmead and Salzberg, 2012) to a custom-modified version of the TAIR10 genome incorporating a specific translocation. Both broad and narrow peaks were called using macs2 v2.2.7.1 (Zhang et al., 2008) in paired-end mode with an input control and a specified genome size. BEDTools intersect v2.26.0 (Quinlan and Hall, 2010) was used to identify reproducible peaks shared across replicates. Coverage files were converted to BigWig format and normalized using an effective genome size of 119,481,543 with deepTools bamCoverage (Ramírez et al., 2014). The normalized coverage for 2 or 4 replicates was averaged using deepTools bigwigAverage.

For differential binding analysis, reads were aligned to the Arabidopsis TAIR10 genome (version 58) using Bowtie2 v2.5.4 (Langmead and Salzberg, 2012). The resulting alignments were converted from SAM to BAM format and sorted using SAMtools v1.21 (Li et al., 2009). Duplicate reads were marked, and multimapped as well as unmapped reads were removed using Sambamba (Tarasov et al., 2015).

Reads overlapping blacklisted regions (Yin et al., 2021) were filtered out using BEDTools intersect (Quinlan and Hall, 2010). Additionally, reads mapping to mitochondrial or chloroplast DNA were removed with SAMtools. Peak calling was performed using MACS2 v2.2.7.1 (Zhang et al., 2008) in paired-end mode with an input control and a specified genome size. Both broad and narrow peaks were identified. Peaks detected in at least two samples were analyzed for differential binding using DiffBind (Stark and Brown, 2011) with DESeq2 option (Love et al., 2014). The results were filtered to retain only those with a false discovery rate (FDR) of less than 0.05. Since only a few differential binding sites were identified, they were manually inspected. Visualization of ChIPseq data was done through the JBrowse platform (Buels et al., 2016).

### DNA isolation and telomere length analysis

Genomic DNA was extracted using the protocol of Dellaporta et al. (1983). For primer extension telomere repeat amplification (PETRA), the Dellaporta protocol was extended with phenol-chloroform-isoamyl alcohol extraction.

Bulk telomere lengths were analyzed using the terminal restriction fragments (TRF) method (Fojtová et al., 2015). In brief, 5 μg of DNA were digested with *MseI* restriction endonuclease (NEB) and separated using 0.8% (w/v) agarose gel electrophoresis. The DNA fragments were transferred to a positively charged nylon membrane (Hybond^TM^ N+, Amersham) and Southern hybridized with a radioactively labeled telomeric probe synthesized by non-template PCR (Adamusová et al., 2020; Ijdo et al., 1991), for sequences of primers see Table S4. The hybridization signals were visualized using a phosphoimager Typhoon FLA 9500 (FujiFilm) and evaluated using the WALTER toolset (Lyčka et al., 2021). The numbers of plants analyzed are given in Table S3.

The lengths of telomeres at individual chromosome arms were assessed using the PETRA method (Heacock et al., 2004). Shortly, a primer extension was performed using 500 ng of DNA and an adaptor primer targeting the G-overhang of telomeres (PETRA-T). Extension products were PCR amplified using a subtelomeric primer specific to the chromosome arm of interest and a primer PETRA-A reflecting the sequence of the tag on the 5′ end of the PETRA-T primer. DMSO was added to increase the annealing specificity of primers. PCR products were analyzed using Southern hybridization with a radioactively labeled telomeric probe; hybridization signals were evaluated using the WALTER toolset. Telomere lengths between combined generations of WTs and generations of mutants were compared by two-tailed Welch’s t-test according to the WALTER algorithm (Lycka et al., 2021). Sequences of primers are provided in Table S4, and the number of plants analyzed is given in Table S3.

## Author contributions

MF, JF and HP provided ideas and the concept of the study. MF and JF provided funding. OH, BM, AH and NB performed experimental work. KH, OH and ML analyzed data, including bioinformatic analyses. MF and OH wrote the manuscript, with input from JF and HP.

## Acknowledgments

This work was supported by the Czech Science Foundation project 22-04364S, and project TowArds Next GENeration Crops, reg. no. CZ.02.01.01/00/22_008/0004581 of the ERDF Programme Johannes Amos Comenius. We thank Jana Kapustová for her excellent technical support. Core Facility Plant Sciences, CEITEC Masaryk University, and Core Facility Genomics, CEITEC Masaryk University supported by the NCMG research infrastructure (LM2018132 funded by the Ministry of Education, Youth and Sports of the Czech Republic) are gratefully acknowledged for their support in obtaining scientific data presented in this paper. Computational resources were provided by the e-INFRA CZ project (ID:90254), supported by the Ministry of Education, Youth and Sports of the Czech Republic.

## Conflict of interest

The authors declare no conflict of interest

## Data availability statement

The author responsible for the distribution of materials integral to the findings presented in this article is Miloslava Fojtová.

## Supporting information

Figure S1. Visualization of RGB-imaging-based parameters as PCA plots and bi-plots.

Figure S2. Telomere lengths analyzed in leaves of T5 plants by the terminal restriction fragments (TRF) method.

Figure S3. Distribution of telomere lengths in WT plants and plants with translocated chromosome arms.

Figure S4: Telomere lengths at individual chromosome arms analyzed in leaves of T5 plants by the primer extension telomere repeat amplification (PETRA) method.

Table S1. Differentially expressed genes.

Table S2. Differential binding analysis.

Table S3. Number of plants analyzed.

Table S4. List of primers.

## Notes

### Competing Interest Statement

The authors have declared no competing interest.

