## Supplementary Figures S1-S4 for "Chromosome engineering points to the *cis*-acting mechanism of chromosome arm-specific telomere length setting and robustness of plant phenotype, chromatin structure and gene expression"

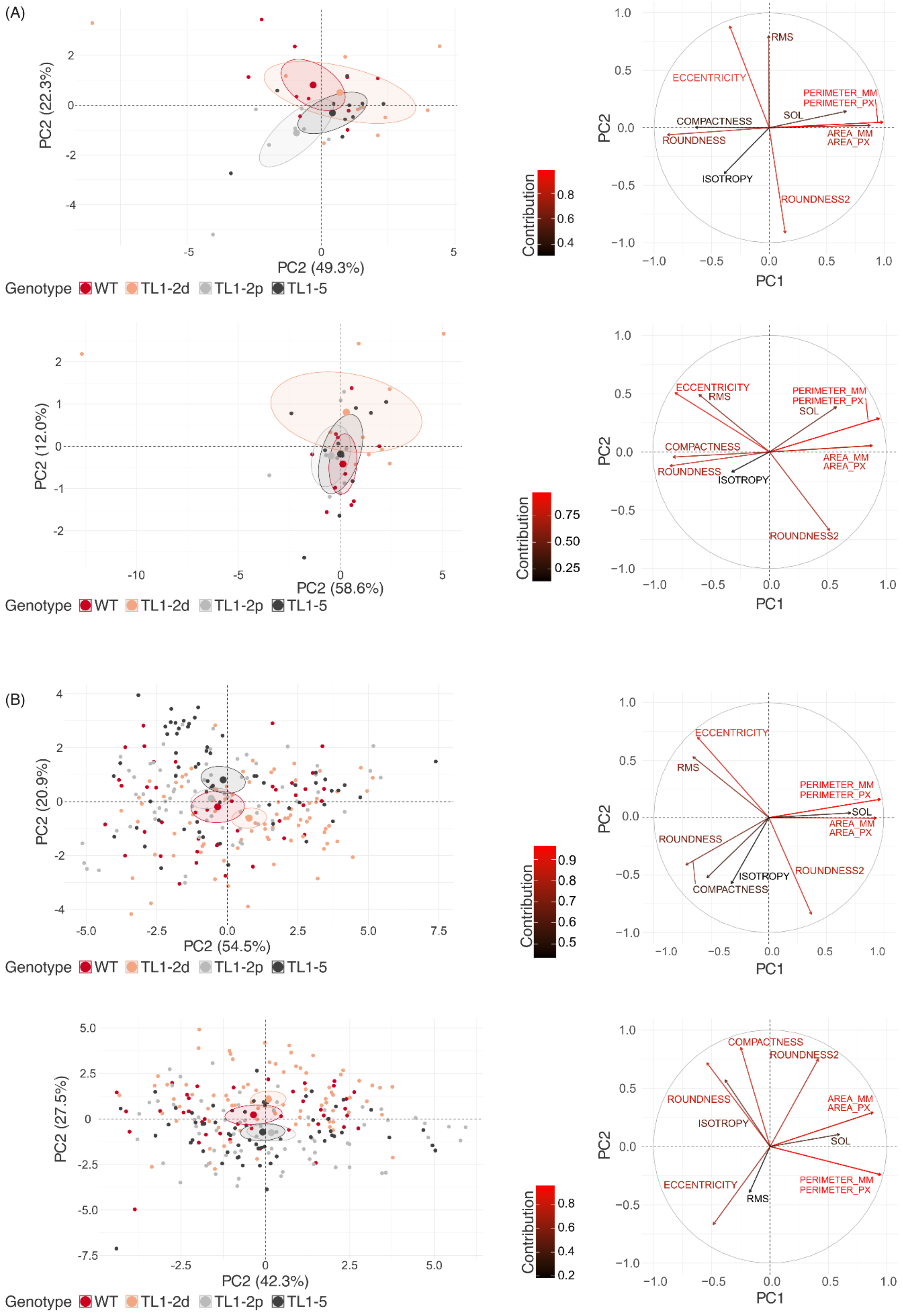

(A)
PC2 (22.3%)
PC2 (49.3%)
Genotype WT TL1-2d TL1-2p TL1-5
Contribution
PC1
PC2
RMS
ECCENTRICITY
PERIMETER_MM
PERIMETER_PX
SOL
COMPACTNESS
ROUNDNESS
AREA_MM
AREA_PX
ISOTROPY
ROUNDNESS2
PC2 (12.0%)
PC2 (58.6%)
(B)
PC2 (20.9%)
PC2 (54.5%)
PC2 (27.5%)
PC2 (42.3%)

Figure S1: Visualization of RGB-imaging-based parameters as PCA plots (left) and bi-plots (right) of plants from T4 (A) and T6 (B) generations at 4 WAS (upper panels) and 5 WAS (lower panels). The length of the arrow (the longer the arrow, the higher the contribution) explains the contribution of tested variables. The differences in plant genotypes are indicated in PCA plots as centroids with ellipses representing their 95% confidence interval.

(A)

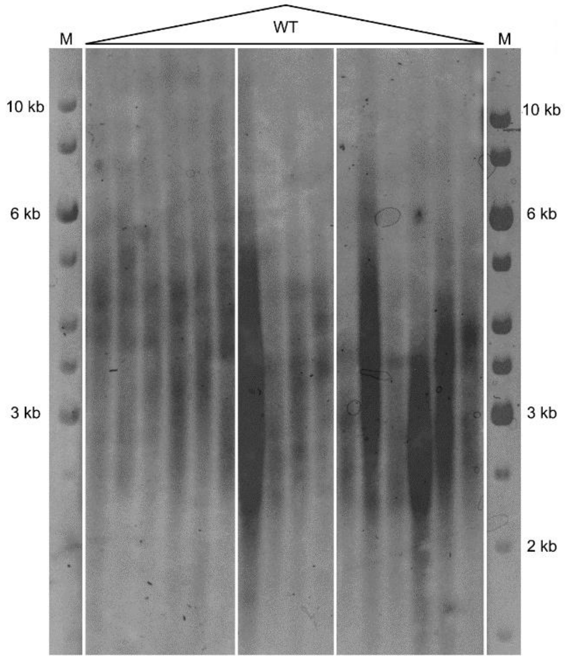
WT
M
M
10 kb
6 kb
3 kb
10 kb
6 kb
3 kb
2 kb

(B)

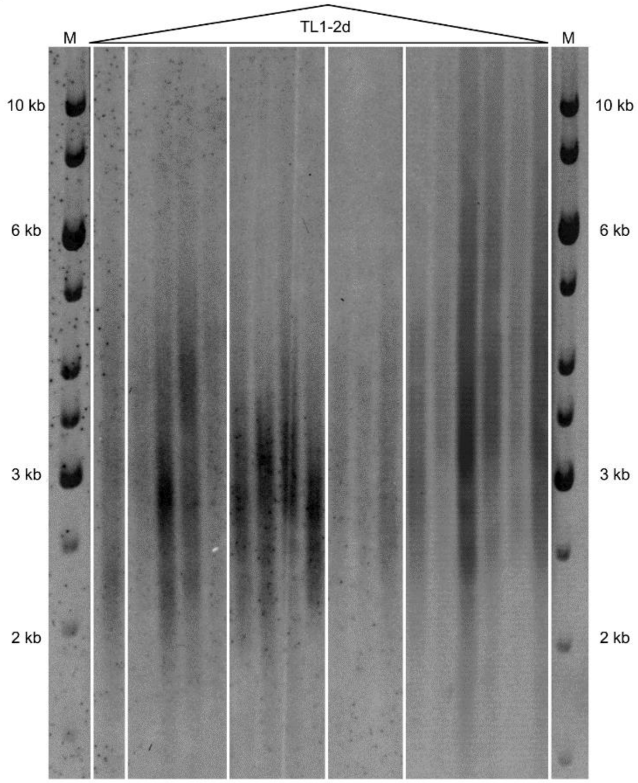
TL1-2d
M
M
10 kb
6 kb
3 kb
2 kb
10 kb
6 kb
3 kb
2 kb

(C)

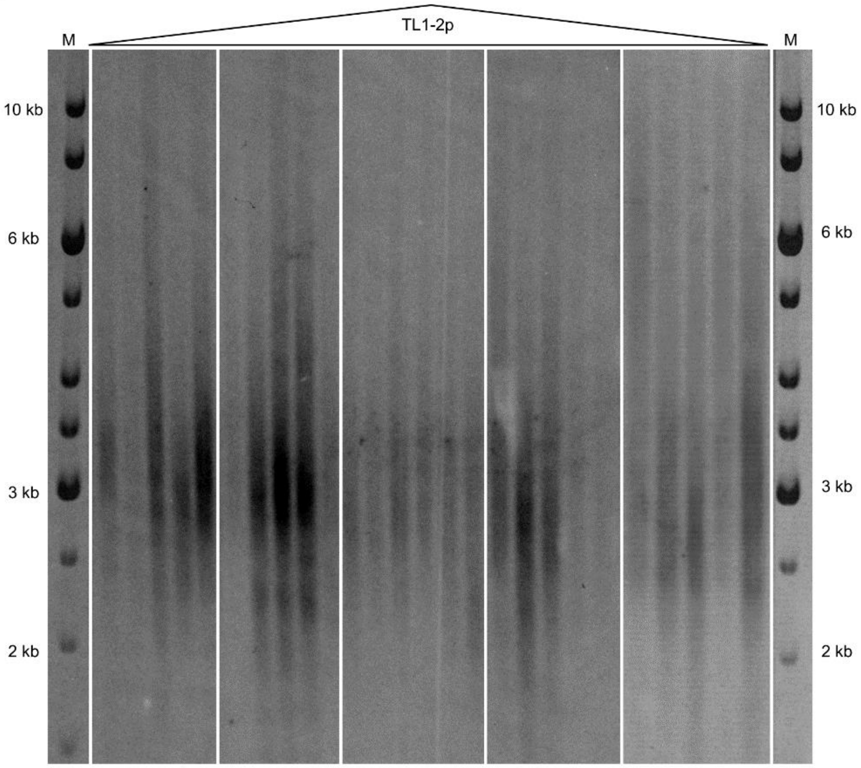
TL1-2p
M
M
10 kb
6 kb
3 kb
2 kb
10 kb
6 kb
3 kb
2 kb

(D)

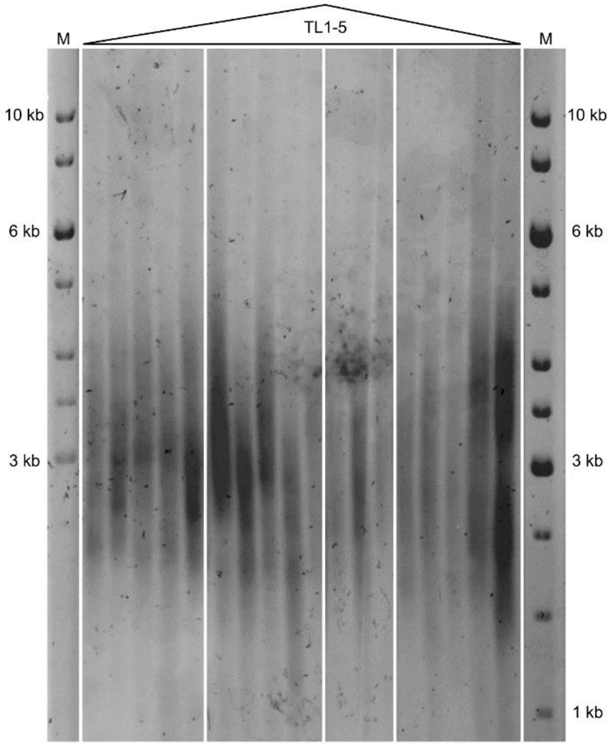
TL1-5
M
M
10 kb
6 kb
3 kb
10 kb
6 kb
3 kb
1 kb

Figure S2: Telomere lengths analyzed in leaves of T5 plants by the terminal restriction fragments (TRF) method. (A) WT plants. (B) Plants with translocated arms of chromosomes 1 and 2 with deletion at the junction site, TL1-2d. (C) Plants with translocated arms of chromosomes 1 and 2 with perfect ligations at both junction sites, TL1-2p. (D) Plants with translocated arms of chromosomes 1 and 5, TL1-5. M, DNA size marker, 1-kb DNA Gene Ruler Ladder, Thermo Fisher Scientific. Telomere-specific hybridization signals were evaluated by the WALTER toolset (Lycka et al., 2021) (Figure S3A).

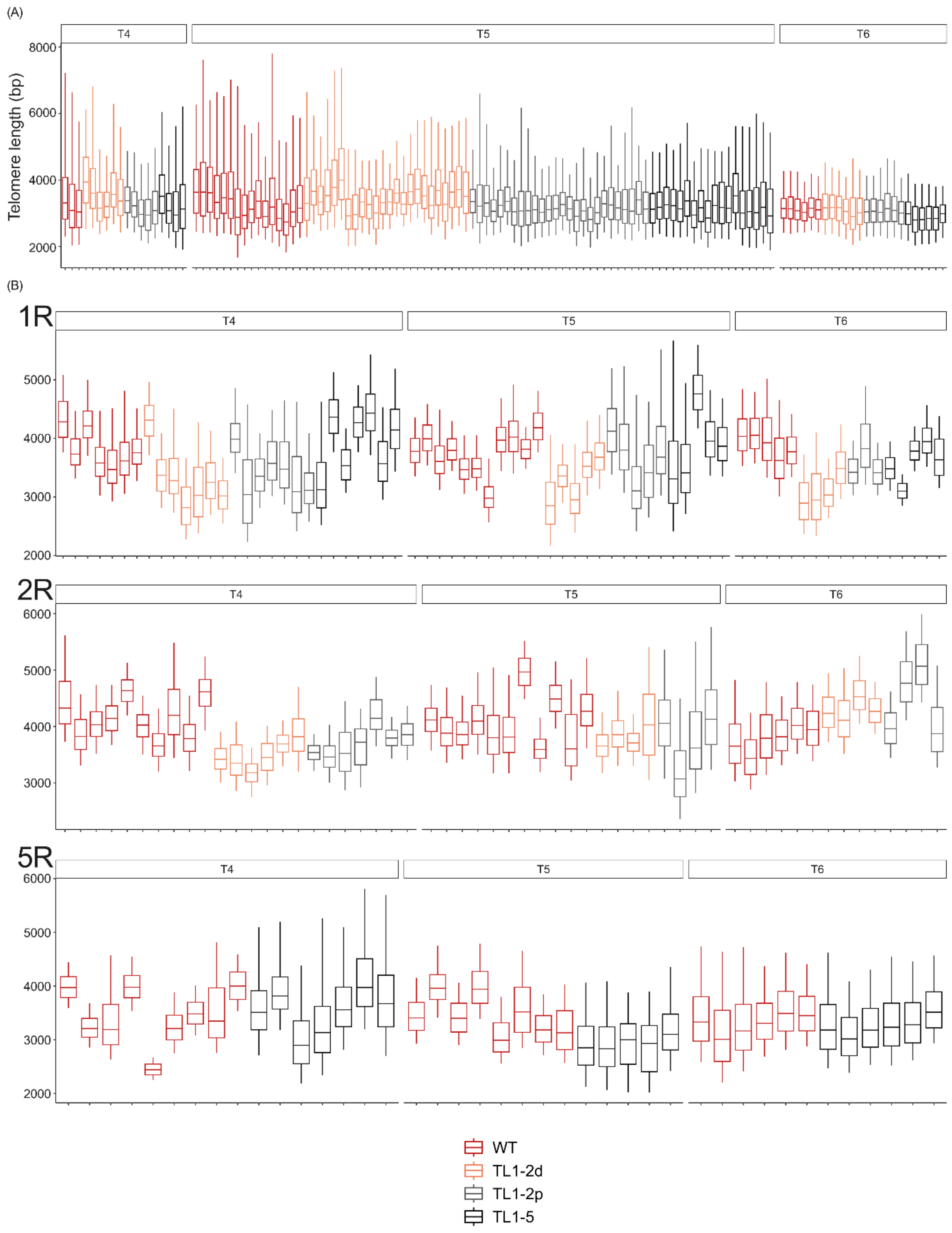
(A)
T4
T5
T6
Telomere length (bp)
8000
6000
4000
2000
(B)
1R
T4
T5
T6
5000
4000
3000
2000
2R
T4
T5
T6
6000
5000
4000
3000
5R
T4
T5
T6
6000
5000
4000
3000
2000
WT
TL1-2d
TL1-2p
TL1-5

Figure S3: Distribution of telomere lengths in WT plants and plants with translocated chromosome arms. Telomere-specific hybridization signals obtained in analyses of telomere lengths by TRF (A) and PETRA (B) protocols were evaluated using the WALTER toolset (Lycka et al., 2021) and presented as boxplots. The central line indicates the weighted median, box limits represent the first and third quartiles and whiskers indicate the minimum and maximum of the selected area of telomeres. (A) Evaluation of hybridization signals of bulk telomeres obtained in analyses of T4, T5 and T6 generations of plants with translocated chromosome arms by the TRF protocol. For raw data of plants of T5 generation, see Figure S2. (B) Evaluation of hybridization signals of individual chromosome arms (1R, 2R, 5R) obtained in analyses of T4, T5 and T6 generations of plants with translocated chromosome arms by the PETRA protocol. For raw data of plants of T5 generation, see Figure S4.

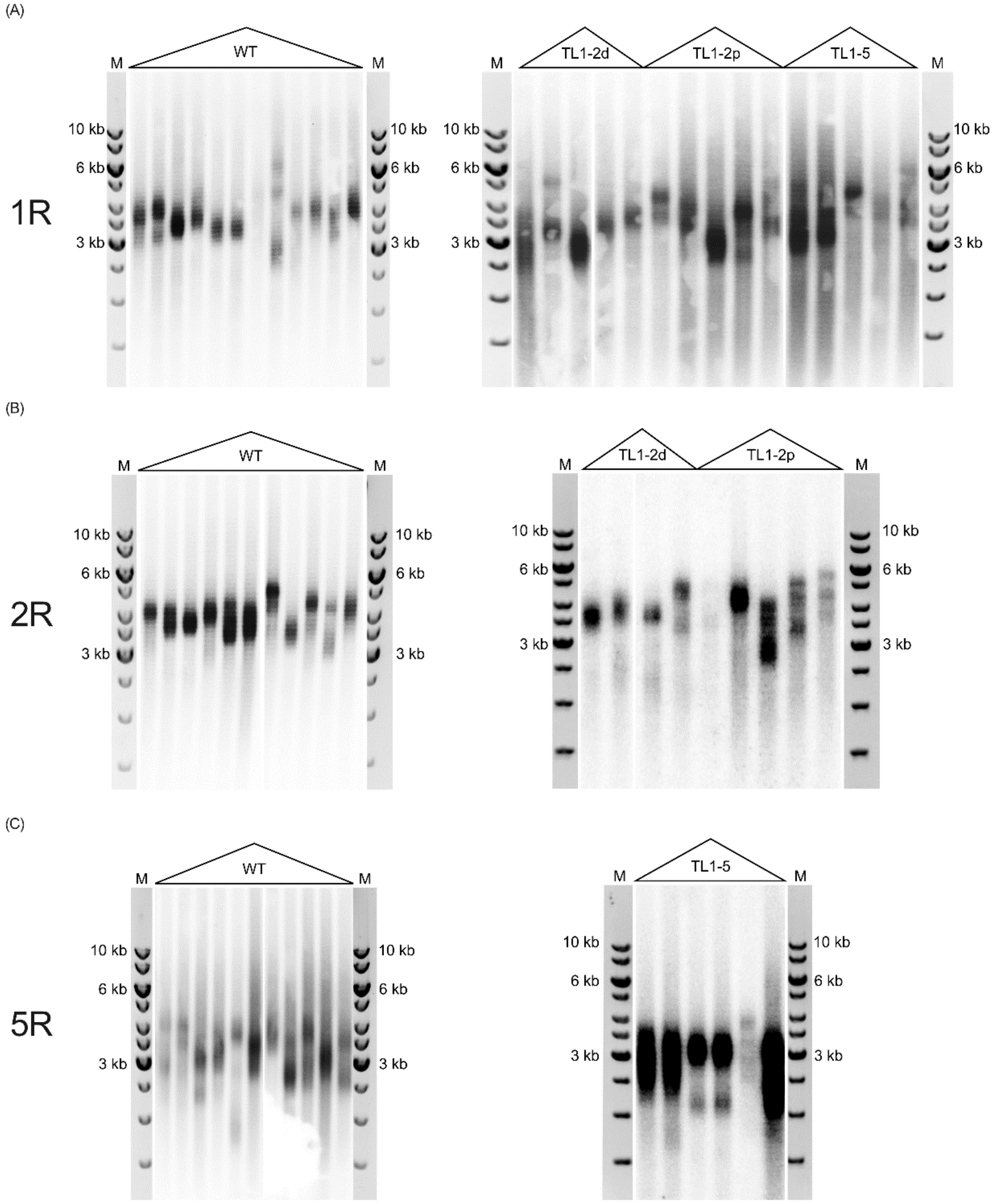


Figure S4: Telomere lengths at individual chromosome arms analyzed in leaves of T5 plants by the primer extension telomere repeat amplification (PETRA) method. (A) Telomeres at the 1R chromosome arms were analyzed in WT plants, plants with translocated arms of chromosomes 1 and 2 with deletion at the junction site (TL1-2d), plants with translocated arms of chromosomes 1 and 2 with perfect ligations at both junction sites (TL1-2p), and plants with translocated arms of chromosomes 1 and 5 (TL1-5). (B) Telomeres at 2R chromosome arms were analyzed in WT, TL1-2d and TL1-2p plants. (C) Telomeres at 5R chromosome arms were analyzed in WT and TL1-5 plants. M, DNA size marker, 1-kb DNA Gene Ruler Ladder, Thermo Fisher Scientific. Telomere-specific hybridization signals were evaluated by the WALTER toolset (Lycka et al., 2021) (Figure S3B).
